# Synthetic DNA templates for the production of *in situ* hybridization probes

**DOI:** 10.1101/108613

**Authors:** B. Rodney Jarvis, Brian G. Condie

## Abstract

Generating RNA riboprobes for *in situ* hybridization generally requires the use of plasmids, which must be grown in bacteria, isolated, purified, and linearized prior to *in vitro* transcription. Here we report a simple method for generating DNA templates for the *in vitro* transcription of RNA probes from synthetic DNA (IDT gBlocks). Each synthetic DNA template contains sequences corresponding to the target mRNA flanked by bacteriophage promoters. Amplification of the template by a single round of PCR and subsequent *in vitro* transcription results in production of high quality RNA probes for *in situ* hybridization.

## Introduction

Typically the generation of RNA *in situ* hybridization probes has been performed using plasmids containing the cDNA of the gene of interest flanked by bacteriophage promoters (Melton et al., 1984). This method requires either PCR amplification of the sequence from a cDNA template and cloning the sequence into a plasmid vector or obtaining a cDNA clone from a colleague or vendor. This is followed by amplification of the cDNA plasmid in bacterial culture and then isolating and purifying the plasmid. The plasmid is linearized with appropriate restriction enzymes and incubated with the appropriate RNA polymerase thereby generating an RNA probe. Sometimes it is preferable to use only a portion of the cDNA clone as a template to generate a probe that is specific to a portion of the mRNA. In those cases a DNA template can be generated from a fragment of the cDNA sequence by performing PCR with primers that flank the desired probe sequence. Typically, the primers contain sequences corresponding to the gene region of interest, as well as sequences of a bacteriophage promoter, such as T7, T3, or SP6. (Kain, Orlandi, & Lanar, 1991; Krieg & Melton, 1987; Wei & Condie, 2011). The PCR product can then be used as a template for the generation of an RNA probe.

These tried and true methods for generating templates to produce RNA probes are costly, labor intensive, and time consuming. However, the development of inexpensive and high fidelity gene synthesis methods provides an approach that avoids many of the technical obstacles in the generation of RNA probes for *in situ* hybridization. As technology has improved, the ability to efficiently and accurately generate synthetic DNA fragments at low cost has increased.

To test the feasibility of using synthetic DNA templates for the generation of RNA probes for *in situ* hybridization we focused on two mouse genes *IL7* and *Foxg1*. We routinely perform *in situ* hybridization to monitor the expression of these genes in various mutant backgrounds. The probes generated from synthetic DNA templates performed extremely well. This approach greatly reduces the time and cost of *in situ* hybridization analyses especially in studies that require the analysis of many genes.

## Methods

Synthetic DNA templates (gBlock Gene Fragments) were ordered from Integrated DNA Technologies, Inc (IDT). Each gBlock contains the sequence corresponding to the mRNA of interest flanked by the T7 and T3 bacteriophage promoters. We incorporated additional sequences at each end of the gBlock to minimize “breathing” of the promoter regions during the transcription reactions.

### *Foxg1* gBlock probe template design

We designed a probe synthesis template for *Foxg1* with the *Foxg1* transcript sequence flanked by T7 and T3 promoters. This template had the following configuration: T7 promoter - *Foxg1* sense strand - T3 promoter. The 5’sequence contained the T7 promoter (*underlined*) as well as 7 random bases preceding the T7 promoter (5’–ATGCATC<u>TAATACGACTCACTATAGGG</u> – 3’). The 3’ end of the fragment contained the anti-sense sequence for the T3 promoter (*underlined)* as well as 7 random bases at the most 3’ end (5’ – <u>CTTTAGTGAGGGTTAATTG</u>TCCATA – 3’). These sequences flanked the cDNA sequence of *Foxg1* from which the anti-sense RNA probe will be transcribed from (5’–GGCACGACCGGCAAGCTGCGGCGCCGCTCCACCACGTCTCGGGCCAAGCTGGCCTTTAAGCGCGGGGCGCGCCTCACCTCCACCGGCCTCACCTTCATGGACCGCGCCGGCTCCCTCTACTGGCCCATGTCGCCCTTCCTGTCCCTGCACCACCCCCGCGCCAGCAGCACTTTGAGTTACAACGGGACCACGTCGGCCTACCCCAGCCACCCCATGCCCTACAGCTCCGTGTTGACTCAAAACTCGCTGGGCAACAACCACTCCTTCTCCACCGCCAACGGGCTGAGTGTGGACCGGCTGGTCAACGGGGAGATCCCGTACGCCACGCACCACCTCACGGCCGCTGCGCTCGCCGCCTCGGTGCCCTGCGGCCTGTCGGTGCCCTGCTCCGGGACCTACTCCCTCAACCCCTGCTCCGTCAACCTGCTCGCGGGCCAGACCAGTTACTTTTTCCCCCACGTCCCGCACCCGTCAATGACTTCGCAGACCAGCACGTCCATGAGCGCCCGGGCCGCGTCCTCCTCTACGTCGCCGCAGGCCCCCTCGACCCTGCCCTGTGAGTCTTTA AGACCCTCTTTGCCAAGTTTTACGACAGGACTGTCCGGGGGACTGTCTGATTATTTCACACAT – 3’). The PCR reaction protocol was as follows; initial denaturation 94 degrees Celsius for 5 minutes, [94°C for 30s, annealing 56°C for 45s, elongation 72°C for 60s] x 33 cycles, final elongation 72°C 10 minutes.

### Interleukin 7 (***IL-7***) gBlock probe template design

We also designed a double stranded cDNA fragment to generate an anti-sense RNA probe to detect the mRNA encoding the cytokine IL-7. This probe template is similar in its design to the *Foxg1* gene fragment with the exception that both the 5’ and 3’ ends contained longer regions of random bases than were used for the *Foxg1* probe template.

(5’- ATGCATCGCGTGCTGCTGGCCTGGCACTG<u>TAATACGACTCACTA TAGGG -</u> (*IL-7* cDNA sense strand sequence) - <u>CCTTTAGTGAGGGTTAATTG</u>TCCAGAGTAGCGGACGGTGACGCCGTCGAGTCAG – 3’). The *IL-7* probe template sequence was flanked by these promoter regions (*IL-7* cDNA sequence 5’ – TCTGCTGCCTGTCACATCATCTGAGTGCCACATTAAAGACAAAGAAGGTAAAGCATATGAGAGTGTACTGATGATCAGCATCGATGAATTGGACAAAATGACAGGAACTGATAGTAATTGCCCGAATAATGAACCAAACTTTTTTAGAAAACATGTATGTGATGATACAAAGGAAGCTGCTTTTCTAAATCGTGCTGCTCGCAAGTTGAAGCAATTTCTTAAAATGAATATCAGTGAAGAATTCAATGTCCACTTACTAACAGTATCACAAGGCACACAAACACTGGTGAACTGCACAAGTAAGGAAGAAAAAAACGTAAAGGAACAGAAAAAGAATGATGCATGTTTCCTAAAGAGACTACTGAGAGAAATAAAAACTTGTTGGAATAAAATTTTGAAGGGCAGTATATAAACAGGACATGTAGTAACAACCTCCAAGAATCTACTGGTTCATATACTTGGAGAGGTTGAAACCCTTCCAGAAGTTCCTGGATGCCTCCTGCTCAAATAAGCCAAGCAGCTGAGAAATCTACAGTGAGGTATGAGATGATGGACACAGAAATGCAGCTGACTGCTGCCGTCAGCATATACATATAAAGATATATCAACTATACAGATTTTTGTAATGCAATCATGTCAACTGC – 3’). The PCR reaction protocol was as follows; initial denaturation 94 degrees Celsius for 5 minutes, [94°C for 30s, annealing 71°C for 30s, elongation 72°C for 60s] x 33 cycles, final elongation 72°C 10 minutes.

### *In vitro* transcription and *in situ* hybridization of sectioned tissue

Digoxigenin (DIG) labeled RNA probes were generated by standard protocols. For the *Foxg1* probe, we used 50% digoxigenin labeled UTP to improve dig-UTP incorporation given the high GC content of the probe sequence. For the *IL-7* probe, we used the standard 35% digoxigenin labeled UTP concentration. Both anti-sense probes were synthesized using the T3 RNA polymerase.

Embryos were collected from pregnant females at different stages of development with noon of the day of plug being embryonic day 0.5 (E0.5). Embryos were then staged more accurately based on somite count or morphological features. Samples were fixed in 4% Paraformaldehyde (PFA) overnight, washed in 1xPBS (DEPC-treated), and dehydrated in a series of washes in 70%, 80%, 90%, 95% Ethanol/DEPC-H2O, and 100% Ethanol. Samples were then paraffin embedded for sectioning by standard procedures and sectioned at a thickness of 12μm. Sectioned samples were collected on glass slides (Fisher-Scientific) and rehydrated. *In situ* hybridization to paraffin-embedded sections was performed as reported previously (Zamisch et al., 2005). Samples were then counterstained in 20% Nuclear Fast Red (NFR).

## Results

The probe generated from the *Foxg1* gBlock^®^ Gene Fragment worked extremely well (Figure 1 A, B). *Foxg1* is expressed in the developing telencephalon and olfactory epithelium and our *in situ* hybridization results recapitulate the expected patterns of expression of this transcription factor at E11.5 (Fig. 1A) (Tao and Lai 1992, Baumer et al. 2002, Kawaguchi et al. 2016). We also detected expression of *Foxg1* at E12.5 in developing thymic epithelial cells (TEC) in the fetal thymus primordium (Fig. 1B) corresponding to the reported expression pattern of this gene (Wei and Condie 2011). The probe generated from the synthetic DNA fragment for *Foxg1* was highly specific which resulted in undetectable levels of background and the samples only required incubation in the chromagen BM-Purple (Roche) for approximately 16-24 hours.

**Figure 1.**
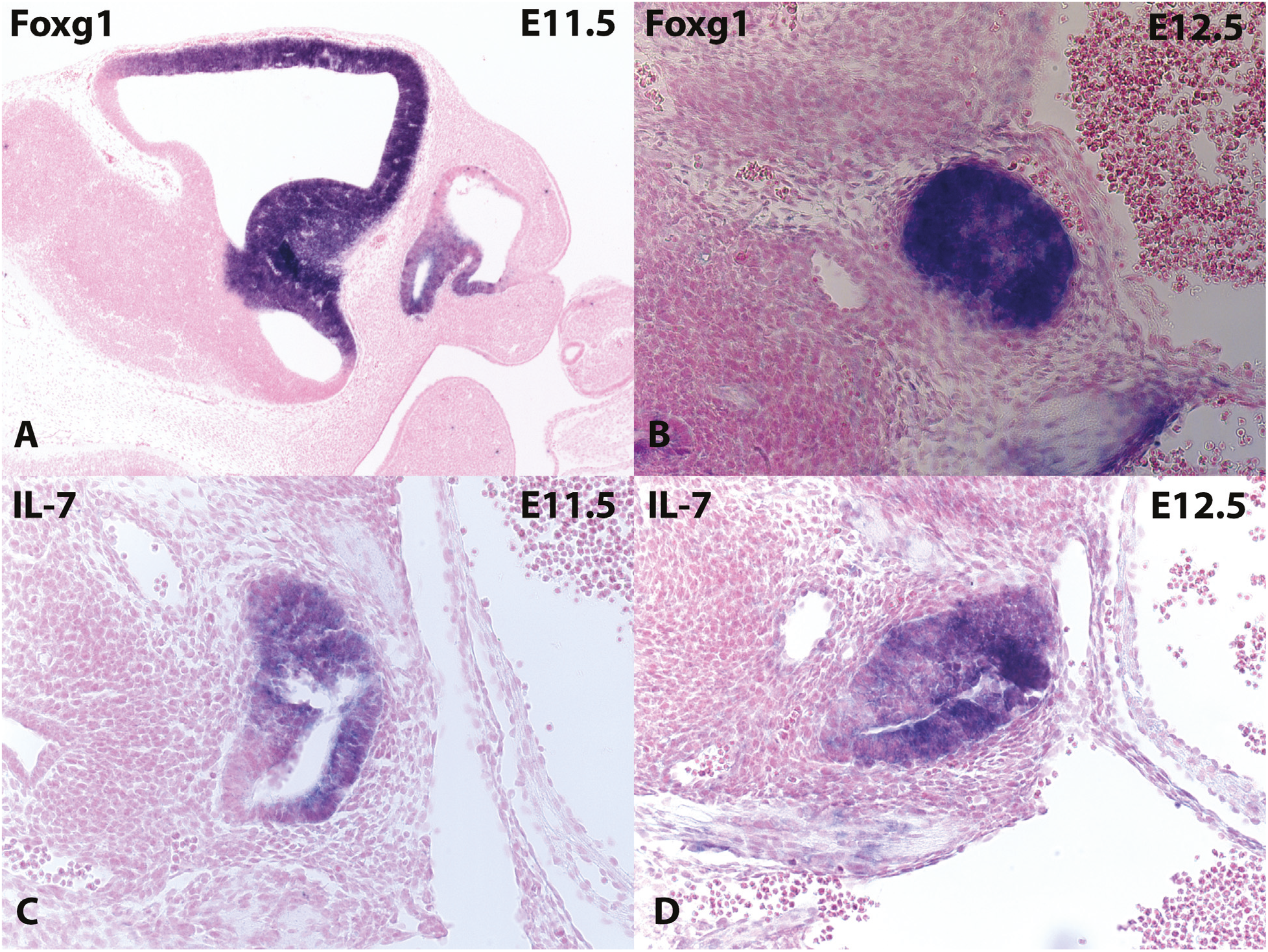
*In situ* hybridization to tissue sections with anti-sense RNA probes generated from synthetic DNA templates. (A, B) Expected patterns of mRNA expression for *Foxg1* in the developing telencephalon at E11.5 (A) and in the thymic primordium (B) at E12.5. (C, D) Expected expression patterns of *IL-7* expression in the thymic primordium at E11.5 (C) and E12.5 (D).

*IL-7* is also expressed in developing thymic epithelial cells where it is responsible for survival and proliferation of developing thymocytes. *In situ* hybridization with our *IL-7* riboprobe mirrored the expression pattern previously reported at E11.5 and E12.5 of mouse development (Fig. 1 C,D) (Zamisch et al., 2005). Sample incubation in BM-purple for this probe required 3-4 days for full development of the chromagen signal. Background was also more variable than the *Foxg1* riboprobe but was still relatively low and comparable to other reported *IL-7* probes (Zamisch et al., 2005).

## Discussion

To optimize the probe template for *IL-7* riboprobe synthesis we had to modify the terminal sequences of the *IL-7* probe gBlock. The *IL-7* gBlock template that we initially designed contained the same terminal sequences as the *Foxg1* gBlock. These terminal sequences were included to provide sites for PCR primers to anneal for amplification of the gBlock and to reduce “breathing” of the template during the transcription reactions. Our intent was to design universal terminal sequences so that we could use the same set of PCR primers for amplification of any gBlock probe template. However, this design resulted in a primer annealing temperature of only 56°C. When we used these primers on the first *IL-7* gBlock design, we repeatedly obtained multiple products. To rectify this issue, we redesigned the *IL-7* template to contain longer primer sequences reported above in the methods. The new primer annealing regions were designed to allow for a Tm of nearly 72°C. This high annealing temperature worked well and eliminated the generation of truncated PCR products.

We have shown that anti-sense RNA probes generated from synthetic DNA templates can perform as well as probes generated from plasmid templates. Overall this approach is simpler, cheaper and faster than plasmid based methods for the generation of RNA probes for *in situ* hybridization. It requires the selection of an appropriate cDNA sequence, flanked by RNA polymerase promoters and with 5’ and 3’ terminal sequences that provide targets for PCR primer annealing. The terminal sequences also reduce “breathing” of the template termini during transcription. This approach greatly simplifies probe generation and reduces costs in the production of large numbers of RNA probes in cases where the expression of many genes must be monitored.

## Literature Cited

Baumer, N., Marquardt, T., Stoykova A., Ashery-Padan R., Chowdhury K., Gruss P. (2002). Pax6 is required for establishing naso-temporal and dorsal characteristics of the optic vesicle. Development 129, 4535–4545.

Kain, K. C., Orlandi, P. A., & Lanar, D. E. (1991). Universal promoter for gene expression without cloning: expression-PCR. Biotechniques 10, 366–374.

Kawaguchi, D., Sahara, S., Zembrzycki, A. and O’Leary D.M. (2016). Generation and analysis of an improved Foxg1-IRES-Cre driver mouse line. Dev. Biol. 412, 139–147.

Krieg, P. A., & Melton, D. A. (1987). In vitro RNA synthesis with SP6 RNA polymerase. Methods Enzymol, 155, 397–415.

Melton, D. A., Krieg, P. A., Rebagliati, M. R., Maniatis, T., Zinn, K., & Green, M. R. (1984). Efficient in vitro synthesis of biologically active RNA and RNA hybridization probes from plasmids containing a bacteriophage SP6 promoter. Nucleic Acids Res. 12, 7035–7056.

Tao, W. and Lai, E. (1992). Telencephalon-restricted expression of BF-1, a new member of the HNF-3/fork head gene family, in the developing rat brain. Neuron 8, 957–966.

Wei, Q., & Condie, B. G. (2011). A focused in situ hybridization screen identifies candidate transcriptional regulators of thymic epithelial cell development and function. PLoS One, 6(11), e26795. doi:10.1371/journal.pone.0026795

Zamisch, M., Moore-Scott, B., Su, D. M., Lucas, P. J., Manley, N., & Richie, E. R. (2005). Ontogeny and regulation of IL-7-expressing thymic epithelial cells. J Immunol, 174(1), 60–67.

